# Multistate structures of the MLL1-WRAD complex bound to H2B-ubiquitinated nucleosome

**DOI:** 10.1101/2022.04.04.486905

**Authors:** Niklas A. Hoffmann, Sanim Rahman, Evan J. Worden, Marissa L. Smith, Kevin E.W. Namitz, Bruce A. Knutson, Michael S. Cosgrove, Cynthia Wolberger

## Abstract

The human Mixed Lineage Leukemia-1 (MLL1) complex orchestrates methylation of histone H3K4 to promote transcription and is stimulated by monoubiquitination of histone H2B. Recent structures of the MLL1-WRAD core complex, which comprises the MLL1 methyltransferase, <u>W</u>DR5, <u>R</u>bBp5, <u>A</u>sh2L, and <u>D</u>PY-30, have revealed variation in the docking of MLL1-WRAD on nucleosomes and left ambiguous portions of Ash2L and the position of DPY30. We used an integrated approach combining cryo-electron microscopy and mass spectrometry-crosslinking to determine structures of the MLL1-WRAD complex bound to ubiquitinated nucleosomes containing the Ash2L intrinsically disordered region (IDR), SPRY insertion region, Sdc1-DPY30 interacting region (SDI-motif), and the DPY30 dimer. We resolved three additional states of MLL1-WRAD lacking one or more subunits, which may reflect different steps in the assembly of MLL1-WRAD. The subunits in all four states are positioned on the nucleosome in manner that is similar to a previous structure of MLL1-WRAD bound to ubiquitinated nucleosome, but that differs from structures with unmodified nucleosomes, suggesting that H2B-ubiquitin favors assembly of the active complex. Our results provide a more complete picture of MLL1-WRAD and the role of ubiquitin in promoting formation of the active methyltransferase complex.

**Significance:** The Mixed Lineage Leukemia-1 (MLL1) complex plays a role in activating transcription by methylating lysine 4 in histone H3, a reaction that is stimulated by the presence of ubiquitin conjugated to histone H2B. Recent structures of the core MLL1 complex, termed MLL1-WRAD, have revealed the existence of multiple docking states and have also left ambiguous portions of the structure. Here we combine mass spectrometry-cross linking with cryo-EM to model additional regions of the MLL1-WRAD complex and identify a series of states that light on complex assembly and the role that ubiquitin plays in orienting MLL1-WRAD on nucleosomes.

## Introduction

Enzymes that deposit post-translational modifications on the core histones, H2A, H2B, H3, and H4, play a central role in regulating transcription in all eukaryotes, from yeast to humans (5). Modifications including acetylation, methylation, and ubiquitination of specific residues impact chromatin structure and recruit enzymes that modulate transcription. Mono-, di-, and tri-methylation of Lys 4 of histone H3 (H3K4) is a hallmark of actively transcribed genes that plays a direct role in transcription activation by stimulating hyperacetylation by the SAGA complex (6,7), initiating the assembly of the transcription pre-initiation complex (8,9), and recruiting nucleosome remodeling enzymes such as CHD1 (10) and NURF (11). H3K4 methylation in most cases depends upon monoubiquitination of histone H2B K120 (H2B-K120Ub) (12-16), which is enriched in actively transcribed regions of the genome (12,17,18).

In humans, H3K4 is methylated by six different methyltransferases from the SET1 family (19), one of which is Mixed Lineage Leukemia-1 (MLL1). The MLL1 catalytic Set domain alone monomethylates H3K4, whereas the enzyme can only di- and trimethylate H3K4 when it is incorporated into the WRAD subcomplex, which also contains WDR5 (tryptophan-aspartate repeat protein-5), RbBP5 (retinoblastoma-binding protein-5), Ash2L (Absent-small-homeotic-2-like) and two copies of DPY-30 (Dumpy-30) (20,21). Chromosomal translocations that result in MLL1 fused to a variety of partner proteins result in MLL-rearranged leukemia (22,23), which has a particularly poor prognosis (24). MLL1 is therefore an attractive therapeutic target, making a better understanding of the molecular basis of MLL1 function inside the WRAD complex important for developing new therapies (25,26).

The Ash2L subunit plays a key role in orchestrating MLL function. Ash2L stimulates MLL1 methylation activity in vitro (27) and knockdown of Ash2L abolishes H3K4 trimethylation in cells (28). Knockout of Ash2L in mice results in embryonic lethality (29). Studies have also shown that transcription factors recruit the WRAD complex by interacting with Ash2L, often in a tissue-specific manner. This includes Ash2L’s reported interaction with the transcription factor, Ap2∂, which recruits the methyltransferase complex to the HoxC8 promoter (30). Ash2L also interacts with TBX1, a transcription factor that is important for heart development and is deleted in human DiGeorge syndrome (29). Mef2d, a transcriptional regulator that targets muscle genes in myoblasts, recruits Ash2L to facilitate terminal differentiation (31). Ash2L has also been shown to shape neocortex formation by transcriptional regulation of Wnt-ß-catenin signaling (32) and plays a key role in maintaining open chromatin in embryonic stem cells (33).

Structural studies have provided insights into the role of Ash2L in MLL methyltransferase complexes. Ash2L is a 60 kDa protein harboring an N-terminal PhD finger, winged helix (WH) DNA binding motif, a SPRY domain (residues 295-484) with a 40-residue insertion termed the SPRY-insertion, and a helical Sdc1-DPY30 interacting region (SDI-motif) (Figure 1a). There is an intrinsically disordered region (IDR, residues 200-295) located between the WH and SPRY domains. Structural and biochemical studies have elucidated the Ash2L WH domain interaction with DNA (34,35). Crystal structures of the Ash2L SPRY domain revealed its beta-sandwich topology (36-38) and interaction with WRAD partners, RbBP5 (38,39) and MLL1 (27,40). The C-terminal SDI-motif of Ash2L protrudes from the SPRY-domain and binds to DPY30 (37,41). A predicted flexible region spanning residues 200-295 (referred to as the IDR (2)) and containing residues important for DNA binding (2), as well as the SPRY-insertion (400-440), were not resolved in the crystal structures.

**Figure 1:**
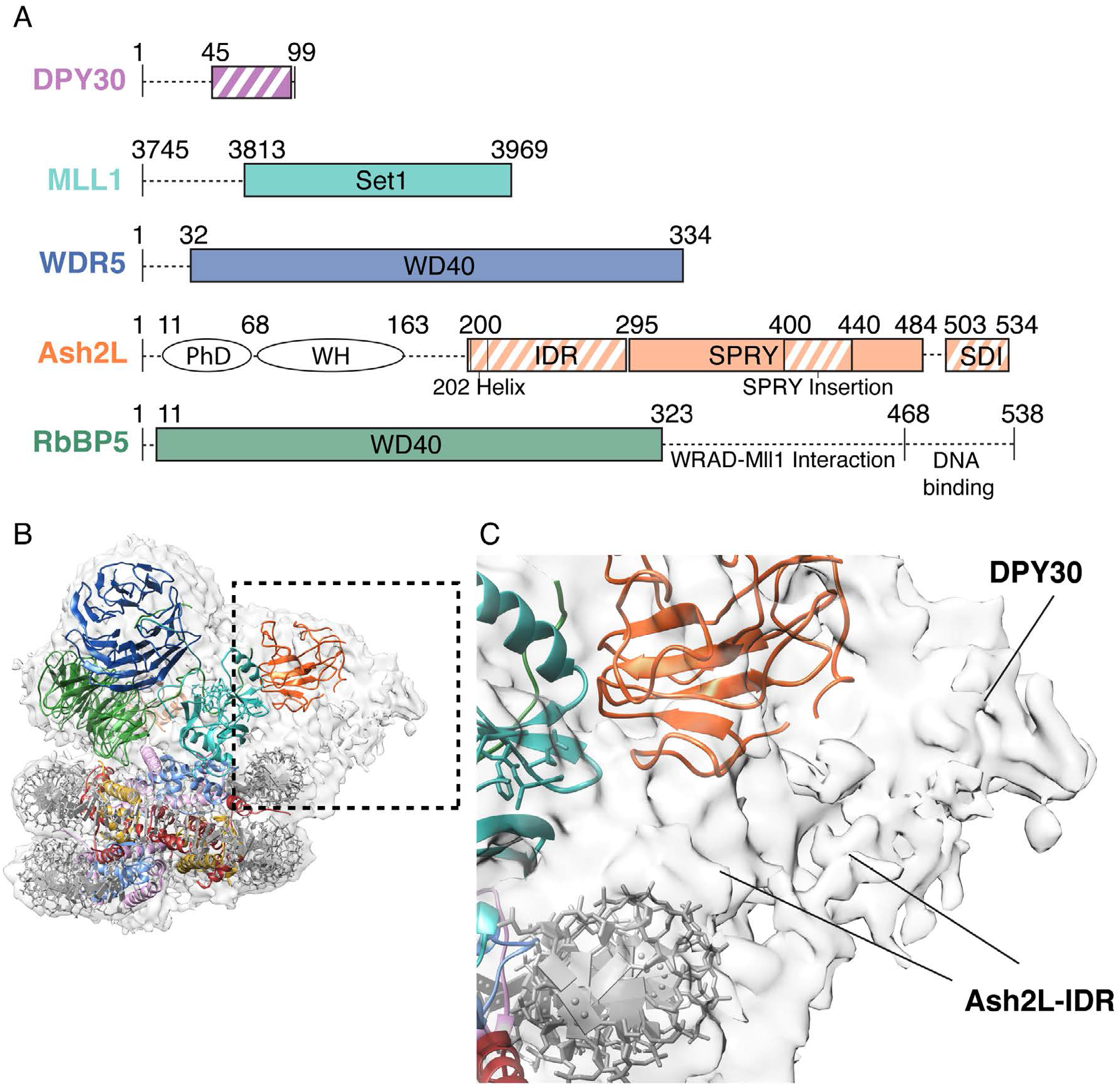
Map of MLL1-WRAD complex shows additional density unexplained by previous structures A) Bar diagram depicting the MLL1-WRAD complex. Circled (not visible in map and PDB) and boxed areas represent structured domains, white-striped boxes show newly added domains in this work. B) Cryo-EM map of MLL1-WRAD calculated at 6 Å resolution showing fit of previously reported structure, PDB 6KIU, superimposed in density. Unaccounted-for density is shown in dashed box. C) Zoom on unaccounted-for density indicating potential subunits that could account for it.

Cryo-EM structures of the MLL1-WRAD complex bound to nucleosomes (1,2) have revealed the overall topology of the complex and its association with the nucleosome. Interestingly, on nucleosomes that lack H2B-Ub, MLL1-WRAD adopts a tilted conformation facing the nucleosome dyad and shows no contacts between MLL1 and the nucleosome (2). In this state, low-resolution density of Ash2L allowed placement of the SPRY-domain, while homology modelling based on the structure of the COMPASS complex, the yeast homologue of MLL1, was used to model the IDR and SPRY-insertions. On ubiquitinated nucleosomes, MLL1-WRAD rotates over the nucleosome disk and adopts what is thought to be an ‘active state,’ with MLL1 contacting the H2A-H2B acidic patch (1). While Ash2L’s SPRY-domain was placed in density in both structures, the IDR and SPRY-insertions are poorly defined and the DPY30 dimer was not placed due to poorly resolved density. In the putative active state structure (1), the authors interpreted weak density between DNA and the SPRY domain as corresponding to an antiparallel beta-sheet of the IDR and the SPRY insertion based on homology to Bre2, the *K. lactis* homologue of Ash2L (42). In the non-active state structure (2), the authors modelled the IDR based on Bre2.

We report here an integrative approach combining cryo-electron microscopy (cryo EM) and mass spectrometry-crosslinking data to model the Ash2L IDR-region and SDI motif in the context of the MLL1 complex bound to H2B-ubiquitinated nucleosomes. Our maps reveal an active state similar to that reported by Xue et al. (1), but with improved density for the region corresponding to the Ash2L IDR and SPRY insertions, as well as DPY30. Using mass spectrometry crosslinking and molecular dynamics flexible fitting (MDFF) (43), we generated a model of the Ash2L IDR and SPRY insertion. We also captured three additional distinct states of the MLL1-WRAD complex bound to ubiquitinated nucleosomes. Together, our results provide new insights into MLL1 structure and assembly.

## Results

### Structure determination of multiple MLL1-WRAD states

We determined the structure of the human MLL1-WRAD complex (21) bound to a ubiquitinated nucleosome using single particle cryo-electron microscopy (cryo-EM). The *Xenopus laevis* nucleosome contained ubiquitin linked to K120 of histone H2B via a non-hydrolyzable dichloroacetone (DCA) linkage (see Methods and (44)). In addition, K4 of histone H3 was substituted with norleucine, a non-natural amino acid which has been shown to bind more tightly to SET methyltransferases than the native lysine (45). Replacing H3K4 with norleucine was previously shown to drive tighter binding of COMPASS, the yeast homologue of MLL1-WRAD, to nucleosomes (46). The minimal fully active human MLL1 fragment containing the catalytic SET domain and spanning residues 3745-3969 was combined with full length RbBP5, Ash2L, DPY30 and WDR5 as previously reported (21) to reconstitute MLL1-WRAD. A complex of MLL-WRAD bound to ubiquitinated nucleosomes was stabilized by crosslinking with glutaraldehyde (see Methods).

The MLL-WRAD complex was imaged with a Titan Krios equipped with a Gatan K3 direct electron detector (Supplementary Table S1). Our approach (see Methods) enabled us to resolve four distinct MLL1-WRAD states bound to the nucleosome at a resolution (FSC 0.143 criterion) of 3.4 Å for State 1, 3.6 Å for state 2, 4.7 Å for State 3 and 4.3 Å resolution for State 4 (Supplemental Figure S1). Subunits from previously reported human MLL1-WRAD structures (1,2) were placed in each map. State 4 fits well with a previously reported MLL1-WRAD structure (PDB ID 6KIU) (1), but also contained additional density adjacent to the Ash2L SPRY domain that was not accounted for by the 6KIU model (Figures 1b and 1c). We hypothesized that this density might correspond to the DPY30 dimer and the more dynamic IDR and SPRY-insertion regions of Ash2L (Figure 1A), which were not resolved in previous reports (1,2). The three additional MLL1-WRAD states identified in our study (Supplemental Figure S1) and described in further detail below, have not been previously reported.

### Crosslinking – mass spectrometry on the MLL1-WRAD complex

To provide an experimental basis for molecular modeling of the extra density in State 4, we used crosslinking-mass spectrometry on the MLL1-WRAD complex to identify close contacts that could be used as restraints in model-building. We crosslinked the MLL1-WRAD complex with BS3, a homobifunctional primary amine reactive cross-linker that has a theoretical maximum crosslinking distance constraint of 34 Å between Cα atoms of crosslinked lysine residues (47). Each MLL1-WRAD subunit contains seven or more lysine residues well distributed throughout the polypeptide, with one or more lysine residues are found in each domain. Cross-linked MLL1-WRAD was digested with trypsin, analyzed by mass spectrometry, and the resulting spectra were searched with the Plink2 cross-link search algorithm (48).

The final dataset comprised 167 lysine-lysine crosslinks at a 2% false discovery rate (FDR) cutoff, with 40 inter-protein crosslinks and 127 intra-protein crosslinks (Figure 2a, Supplementary Table S2). The diverse network of inter-protein crosslinks reflects the intertwined subunit architecture of the MLL1 complex. An exception is DPY30, which only has four cross-links to four residues in Ash2L, consistent with the position of DPY30 at the periphery of the MLL1-WRAD complex. Ash2L is the subunit with the largest number of crosslinks, with 75 inter- and intra-protein crosslinks (Figure 2a). There were 26 inter-subunit crosslinks for the MLL1-Set domain, reflecting this protein’s central position in the MLL1-WRAD complex. By contrast, there were far fewer crosslinks to WDR5 and RbBP5, which are larger proteins and contain a greater number of lysines compared to the MLL1-SET domain. The small number of cross-links to these subunits could be due to the relative inaccessibility of lysines in the WD40 folds of both proteins, which reduces crosslinking efficiency (49). Alternatively, it is possible that there is greater mobility in these regions, which could similarly reduce the number of observed cross-links.

**Figure 2:**
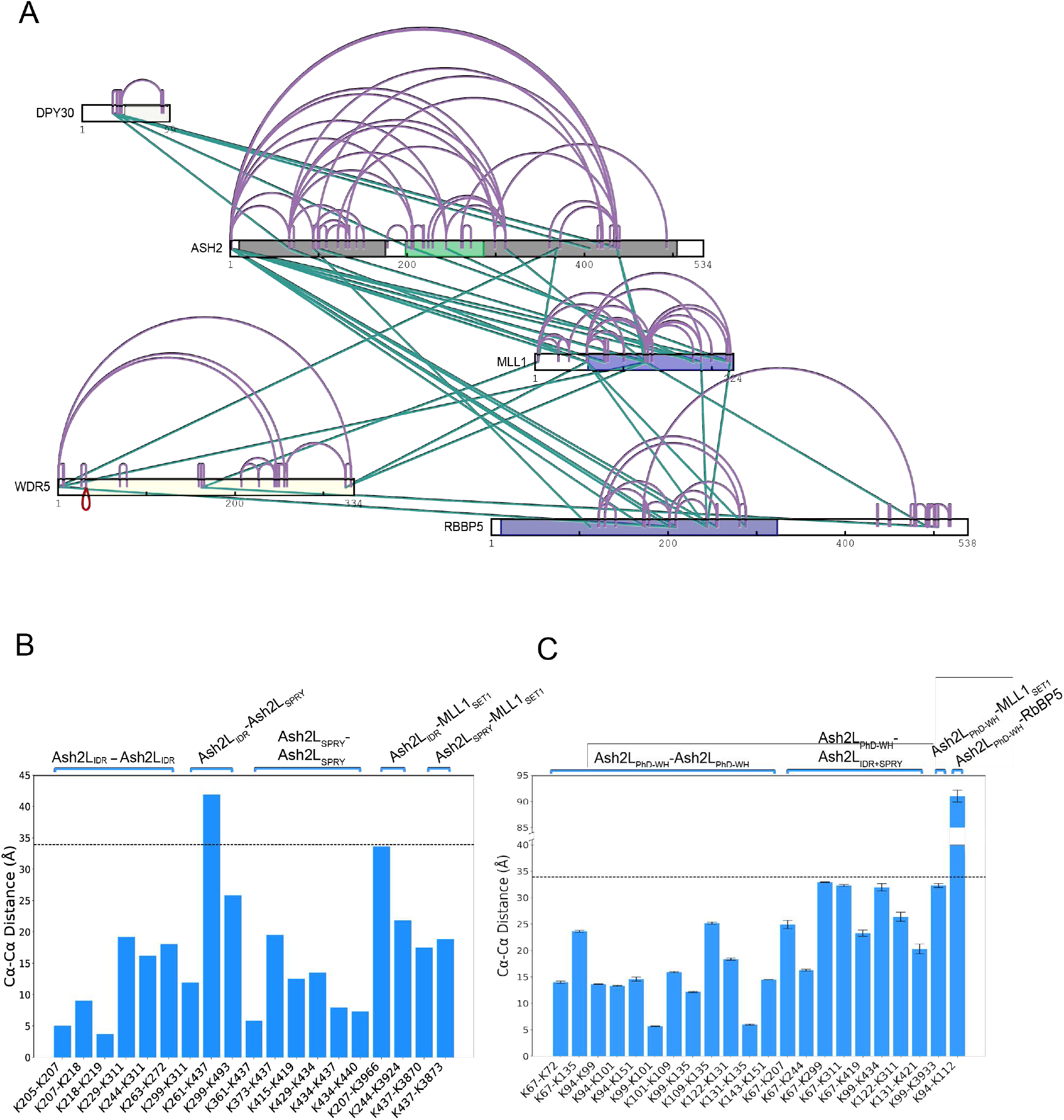
Crosslinks of MLL1-WRAD in solution. A) Intra-protein crosslinks (purple) and inter-protein crosslinks (green) are indicated by lines. B) Calculated Cα-Cα distances between crosslinked lysines in the MLL1_WRAD_-nucleosome complex predicted by HADDOCK involving Ash2L_200-537_. The dashed line denotes BS3’s theoretical maximum crosslinking distance of 34 Å. B) Distances as in (A) calculated for Ash2L_PhD-WH_.

### The DPY30-SDI motif occupies a peripheral density stretch

To fit missing portions of the model to the extra density in our maps, we first utilized two DPY30 crystal structures (PDB 4RIQ and 6E2H) (37,41) and fit them to the density map using the crosslinking data as a guide (see Methods). Both crystal structures contain DPY30 bound to the Ash2L C-terminal SDI motif, which protrudes from the Ash2L SPRY domain (37). This portion of human WRAD was either absent or poorly defined in previous structures of the MLL1 complex bound to nucleosomes (1,2). By placing the SPRY domain of crystal structure 6E2H (37) in our map, the Ash2L SDI motif (residues 503-523) and DPY30 from the crystal structure were near the unallocated density (Figure 3c). A simple rotation of the SDI-DPY30 portion of PDB 6E2H relative to its SPRY domain positions these domains in the density (Figure 3, Supplemental Figure S3b). This placement is supported by unbiased fitting of the SDI-DPY30 substructure to our map using the Fit-in-Map function in Chimera (50). The 15-residue linker (residues 485-500) connecting the Ash2L SDI-DPY30 region and the Ash2L SPRY domain was manually rebuilt and then refined using Molecular Dynamics Flexible Fitting (MDFF) (see Methods). The resulting position of DPY30 and the Ash2L SDI motif fits well into the active state map and satisfies a crosslink (SPRY-K299 to SDI-Linker-K493) (Figures 2b), consistent with the conformational rearrangement modeled here.

**Figure 3:**
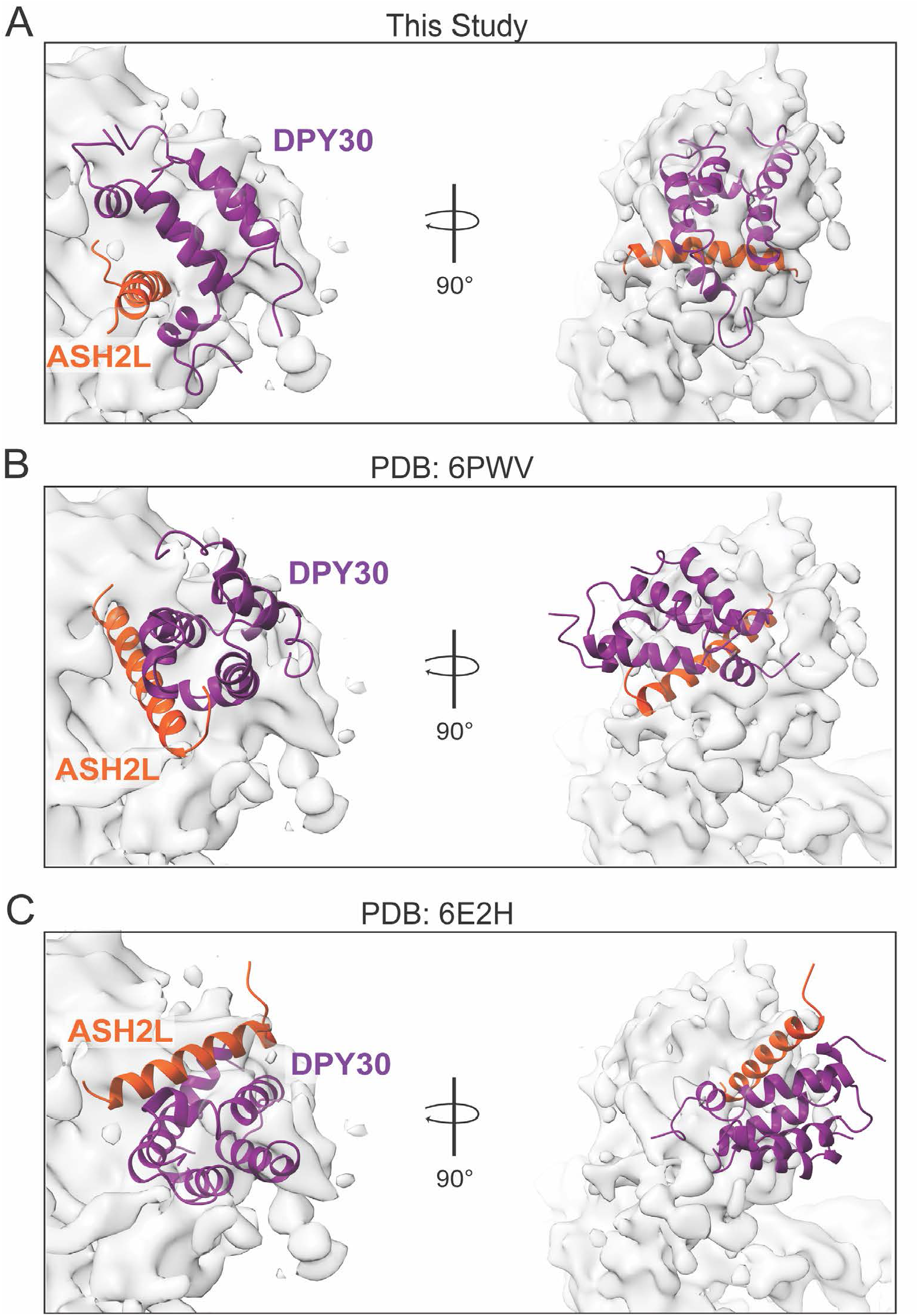
Comparison of Ash2L-SDI motif (orange) and DPY30 dimer (purple) positioning on active state map. All structures were compared by aligning the Ash2L-SPRY domain. A) Structure of the Ash2L-SDI motif and DPY30 dimer presented in this study. B) Structure of the Ash2L-SDI motif and DPY30 dimer reported by Park et al. (6PWV) positioned in our active state map. C) Crystal structure of the Ash2L-SDI motif and DPY30 dimer reported by Haddad et al. (6E2H) positioned in our active state map.

The ability of the Ash2L linker and the SDI-motif protruding from the SPRY-domain to adopt multiple conformations had been observed in previous structures (37,41) (Figure 3, Supplemental Figure S3b). In our model, the partially bent SDI-motif most closely resembles the helix conformation observed in the crystal structure of DPY30 alone bound to the SDI motif (4RIQ) (41) but differs from that previously reported for the cryo-EM structure of MLL1 bound to a nucleosome, 6PWV (Figure 3b, Supplemental Figure S3b). We note that orientation of the SDI motif in our structure is dictated by the positioning of the DPY30 dimer, which is bound to the SDI motif. Whereas the reported position of the SDI-motif in 6PWV could be fit in density in our active state map (Figure 3b, Supplemental Figure S3b), the resulting position of the bound DPY30 dimer does not fit the density in our map (Figure 3b, Supplemental Figure S3c). In further support of our positioning of DPY30, the location of the DPY30 dimer on the complex periphery is consistent with the absence of cross-links between DPY30 and any other subunit except for Ash2L (Figure 2a). The position of DPY30 is also consistent with observed cross-links between DPY30 residues K35 and K40, which are located in a disordered region, to surface lysine K406 and K440 of the Ash2L SPRY domain (Supplementary Table S2).

### Integrative modelling of Ash2L

Placement of DPY30 and the Ash2L SDI motif left unassigned density adjacent to Ash2L that could potentially correspond to the IDR region (residues 200-300) and the SPRY insertion (residues 400-440) (Figure 1A). There is no high-resolution structural information on the Ash2L IDR and previously proposed models of this region based on the Ash2L yeast homologue, Bre2 (2) do not fit our EM map well, nor does a recently reported model generated by AlphaFold (51). The 40-residue insertion in the Ash2L SPRY domain has been shown to contact DNA (1,2) and a homologous stretch of yeast Bre2 similarly extends towards nucleosomal DNA in COMPASS (52,53). We used these observations, combined with cross-linking restraints and MDFF, to model the IDR and SPRY insertion domain in our map (Figure 4b).

**Figure 4:**
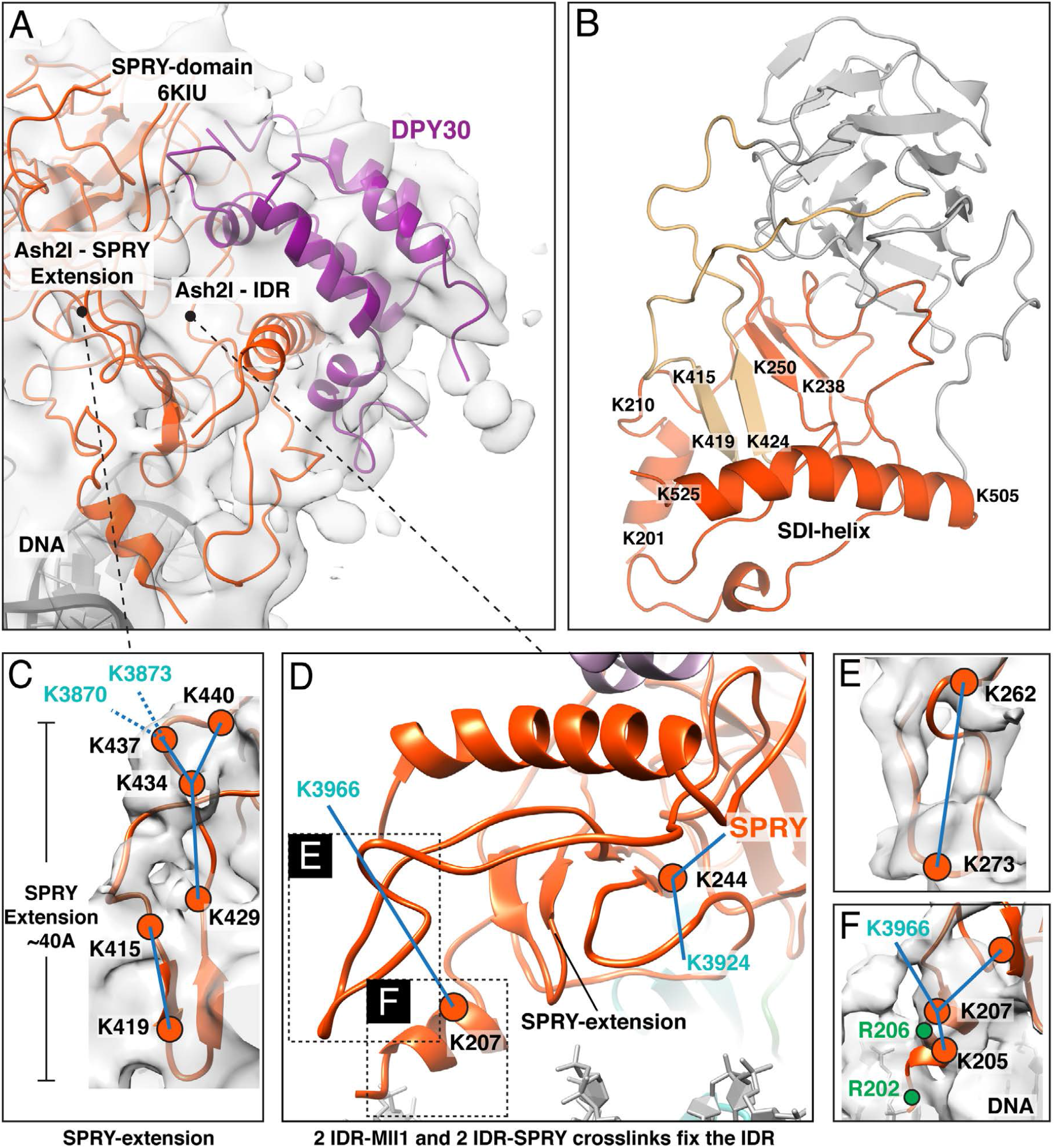
Building and fitting of the Ash2L_IDR-SPRY_ domain to experimental density. A) Placement of Ash2L_IDR-SPRY_ and DPY30 model into the active state map. B) Overview of the Ash2L_IDR-SPRY_ model built by integrative modeling and MDFF refinement. The IDR is colored orange and the SPRY-insertion region is colored in gold. Regions colored in gray correspond to the SPRY domain, which was built using 6KIU as a reference model. C) SPRY-insertion stretching over 40Å fitted to EM-density. Obtained crosslinks are displayed (blue – satisfied). Residues labeled in cyan refer to satisified crosslinks to the MLL1 SET domain. D) IDR-region located in the vicinity of the SPRY-insertion. Intra-SPRY crosslinks from K207 and K244 are displayed and demonstrate the anchoring of the IDR-region. E) Zoom on indicated inset in panel (D) showing IDR-loop 254-270 and satisfied crosslink. F) Zoom on indicated inset in panel (D) showing IDR-helix 200-210, all crosslinks are shown, lysine residues are depicted in orange, arginine residues in green.

The resulting model of Ash2L places the 40-residue SPRY-insertion loop in the EM density at the periphery of the SPRY domain (Figure 4c). The loop extends over 40 Å on the periphery of the SPRY domain to the nucleosomal DNA and contains an anti-parallel beta sheet comprising residues 415-419 and 424-428 (Figure 4c). This positions K419 and K421 near the DNA, consistent with the deleterious effect of substitution of these residues on DPY30-dependent methylation by MLL1 (3). This fit satisfies eight crosslinks, including two inter-subunit crosslinks between Ash2L residue K437 and MLL1 residues K3870 and 3873 (Figure 4c). As a next step, we placed IDR residues K207 and K244 in the unassigned EM density based on crosslinks between these residues with multiple lysine residues within Ash2L, as well as to MLL1 (Figure 4d). Specifically, Ash2L K207 crosslinks to two adjacent IDR lysines (K205, K218) and to MLL1 K3966 (Figure 4d and 4f), while Ash2L K244 crosslinks to the SPRY-domain K311 and to MLL1 residue K3924 (Figure 4d). Next, we included previously reported secondary structure analysis and Bre2 homology that predicts beta-strand features in residues 238-253 (1,2) to extend this region from lysine 244. This stretch comprises two antiparallel beta strands, 238-242 and 247-250, that associate with the two beta strands in the SPRY insertion to form a four-stranded antiparallel beta sheet (Figure 4b). Based on secondary structure prediction (Supplementary Figure S4a), we modeled Ash2L residues 200-210 as an α-helix and placed it in corresponding density (Figure 4f). Notably, the proximity of the helix to DNA (Figure 4a) is consistent with the observation that alanine substitutions of K205/K206/K207 markedly reduce MLL1 methyltransferase activity on nucleosomes (2). Finally, we fit Ash2L residues 253-278 to a short stretch of unassigned density extending to the periphery of the complex (Figures 4d and 4e), and in a position that satisfied the observed crosslink between K262 and K273 (Figure 4e, Supplemental Figure S4b).

The proximity of the Ash2L IDR and SPRY-insertion region to nucleosomal DNA in our model (Figure 4a), could account for the importance of DPY30 binding to MLL1-WRAD activity. Previous studies have shown that DPY30 greatly increases MLL1 methyltransferase activity on nucleosomes and that this increase in activity depends upon the Ash2L IDR (3). DPY30 forms multiple contacts with the IDR and the SDI helix, which help to stabilize folding of the IDR, as supported by NMR studies(3). This ordering of the IDR, in turn, would favor the predicted interactions with nucleosomal DNA, thus accounting for the importance of these residues to MLL1 activity on nucleosomes (2).

There was no apparent density corresponding to the Ash2L N-terminal winged helix (WH) domain (residues 11-67), which has been shown to bind DNA (34,35), nor was there density for the atypical PHD finger (residues 68-163). Previous cryo-EM studies of MLL complexes also did not locate these domains (1,2). We therefore utilized the crosslinks we identified involving the Ash2L N-terminal WH and PHD domains as restraints to guide docking of the crystal structure of these domains (PDB ID 3RSN) (34) to our active state model (Figure 5a). Of the 22 crosslinks we obtained (Supplementary Table S2), the twelve crosslinks between lysines within the WH and PHD finger domains and were satisfied by the crystal structure, indicating that this fragment of Ash2L did not change conformation. The additional ten crosslinks were used as restraints to dock the crystal structure on our MLL1-WRAD active state model. Eight of the crosslinks connected to other regions of Ash2L (Figures 5b, 5c) and two crosslinks connect to MLL1 (Figure 5d) and RbBP5 (Figure 2c and Supplementary Table S2). In the highest scoring model, all crosslinks are satisfied except for one connecting K94 of the Ash2L WH and PHD finger domains to K112 of RbBP5 (Figure 2c) and the top four hits position the WH and PHD finger comparably, with a backbone RMSD of 0.336 ± 0.045 Å. The modeled position of the Ash2L WH and PHD finger domains (Figure 5a) lies in a groove between MLL1 and the Ash2L SPRY domain. Importantly, the model places two lysine residues, K131 and K135, in contact the sugar-phosphate backbone of the nucleosomal DNA (Figure 5e), consistent with their importance to binding of Ash2L to DNA (34). Given the positioning of the WH at the end of the DNA in the nucleosome core particle, it is quite possible that this domain would make additional contacts with DNA that extends beyond this minimal fragment.

**Figure 5:**
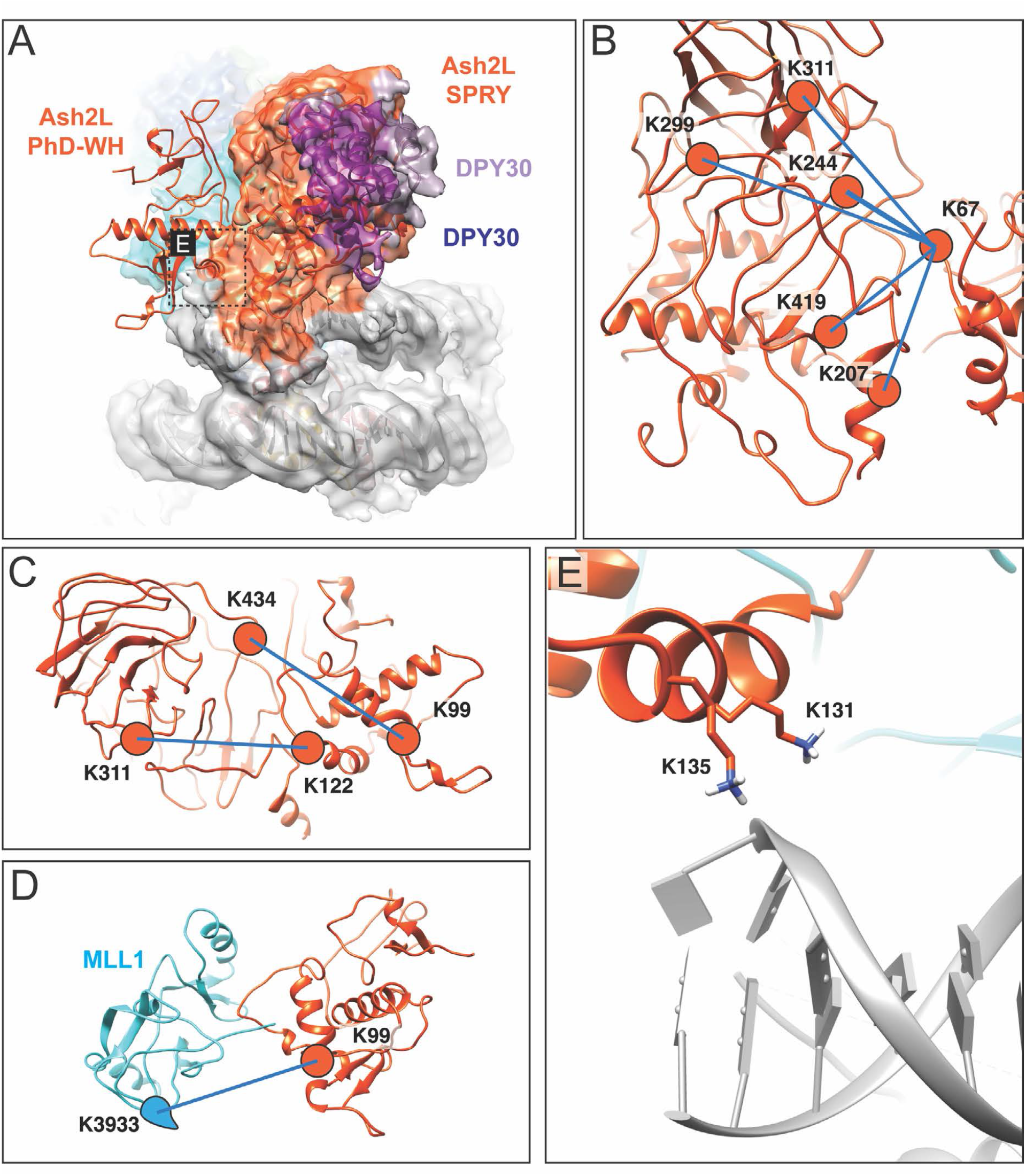
Modelling of Ash2L PHD-WH domain (**A**) Model of Ash2L PhD-WH ensemble on the active state model predicted by crosslinked restrained docking. (**B**) Crosslinks between Ash2L PhD-WH residue K67 and the Ash2L IDR and SPRY domain. (**C**) Additional crosslinks between K99 and K122 in the Ash2L PhD-WH with K311 and K434 in the SPRY domain. (**D**) Single crosslink connecting Ash2L PhD-WH residue K99 to MLL1 SET domain residue K3933. (**E**) Zoom on indicated inset in panel (A) showing the positioning the K131 and K135 in close proximity to the backbone of the nucleosomal DNA.

### Four MLL1-WRAD states represent putative assembly intermediates

In addition to the active state MLL1-WRAD complex described above, we were able to resolve three additional states in our data set (Figure 6, Supplementary Figure S1). All four states have well-resolved density for RbBP5-WD40, whereas other MLL1-WRAD subunits are less well resolved but seem to be in similar positions as compared to the active state model. In State 1 (Figure 6a), only the RbBP5-WD40 domain shows strong density on the nucleosome. There is no density corresponding to remainder of Rbpb5 or to the methyltransferase subunit, MLL1. In state 2 (Figure 6b), WDR5 is also well-resolved, in addition to strong RbBP5-WD40 density, while the remaining MLL1-WRAD density is poorly resolved. State 2 contains additional strong density contiguous with the DNA at SHL-6 to SHL-7 (Figure 6b) that may correspond to a slight peeling away of the DNA from the histone core, although the density is not sufficiently well-resolved to rule out alternative models that place additional protein in this location. State 3 contains equally well resolved density for RbBP5-WD40 and WDR5 as well as better-resolved MLL1-SET domain density and low-resolution Ash2L density (Figure 6c). Interestingly, in this state, the DNA density at SHL-6 to SHL-7 is very weak, as is the MLL1-density as it approaches the nucleosome disk.

**Figure 6:**
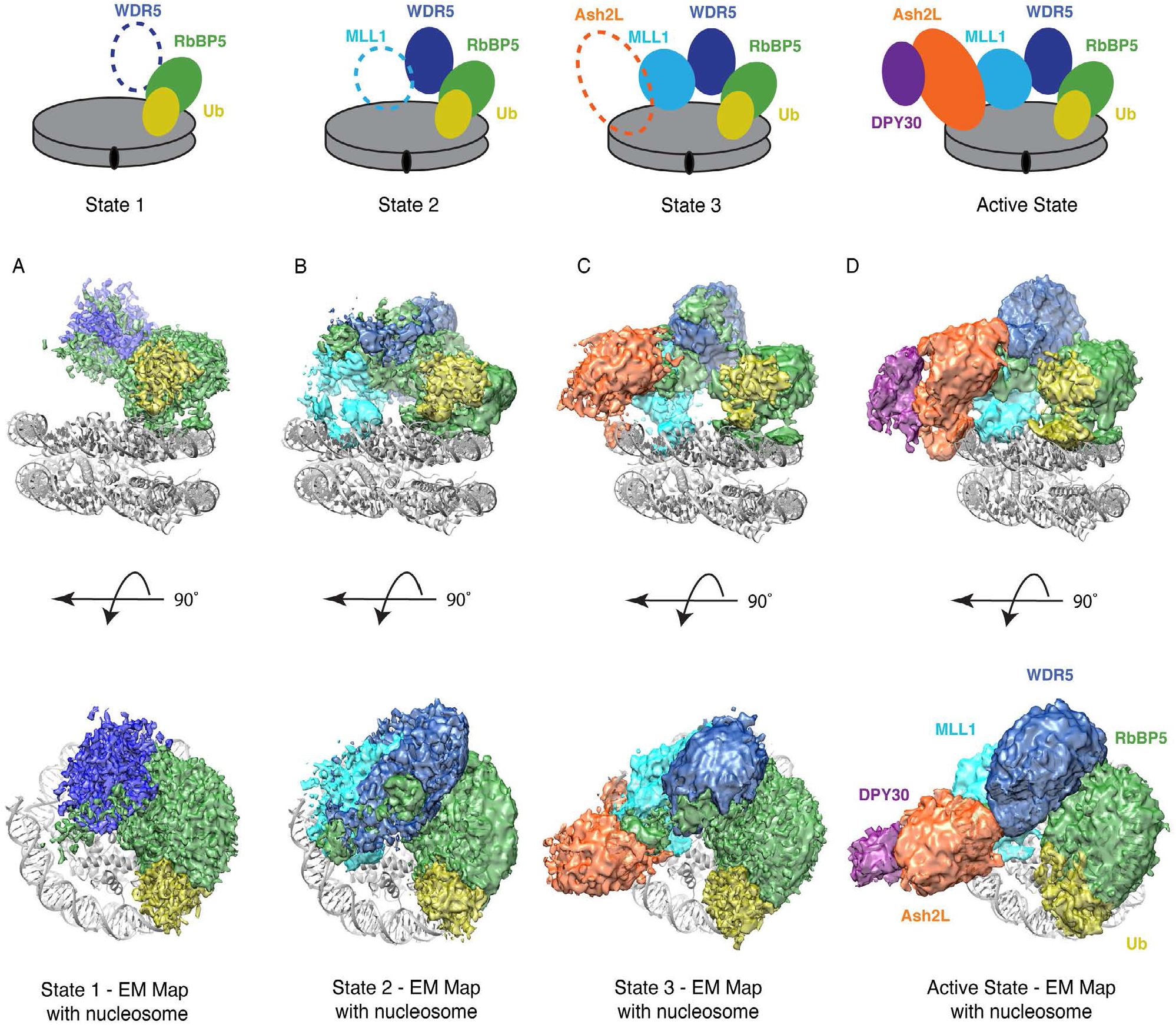
Four distinct states shown stabilization of MLL1-WRAD (define) on the nucleosome. EM maps of the 4 states are shown, with the structure of the active state model superimposed on each map. A) State 1, B) State 2, C) State 3, D) Active state map calculated at 6 Å, shown for comparison. Densities in all panels colored according to subunits as shown in panel D.

A comparison of the four states reveals no apparent change in the docking of the MLL1-WRAD complex on the nucleosome, but rather an apparent progressive assembly of subunits from States 1-3 (Figure 6a-c) and, finally, the active state (State 4) containing all five subunits (Figure 6d). While it is not possible to distinguish ordered assembly from simple heterogeneity of states bound to the nucleosome, one possibility is that the RbBP5 WD40 domain forms an anchor on ubiquitinated nucleosomes. RbBP5 binds to the ubiquitin that is conjugated to histone H2B-K120. The bound RbBP5-WD40 could either facilitate stepwise assembly of the full MLL1-WRAD complex on nucleosomes or, alternatively, stabilize the positions of the remaining subunits on the nucleosome. The flexible C-terminal loops of RbBP5 (residues 330–475) contact all MLL1-WRAD subunits except DPY30 (diagrammed in Supplemental Figure S5). We speculate that the RbBP5 C-terminal residues might play a role in the assembly of MLL1’s active state on the nucleosome. Consistent with our model for step-wise assembly of the complex, a hydrodynamic analysis of the MLL1-WRAD complex identified multiple assembly states in the absence of nucleosomes, which included several different substates in addition to the full five-component complex (54).

### Role of ubiquitin and Ash2L/DPY30 in positioning the MLL1 complex on nucleosomes

A striking feature of reported structures of MLL1 complexes (1-3,55) is their highly variable positioning on the face of the nucleosome. While the spatial relation among the five subunits of MLL1-WRAD complex varies very little, the complex has been observed to dock in multiple orientations on the face of the nucleosome in these structures. These orientations can be categorized according to the superhelical positions on the nucleosome of Rbpb5 and Ash2L/DPY30, which define the two ends of the complex. The positions of these subunits in the present and in previous studies are listed in Table 1.

**Table 1.**
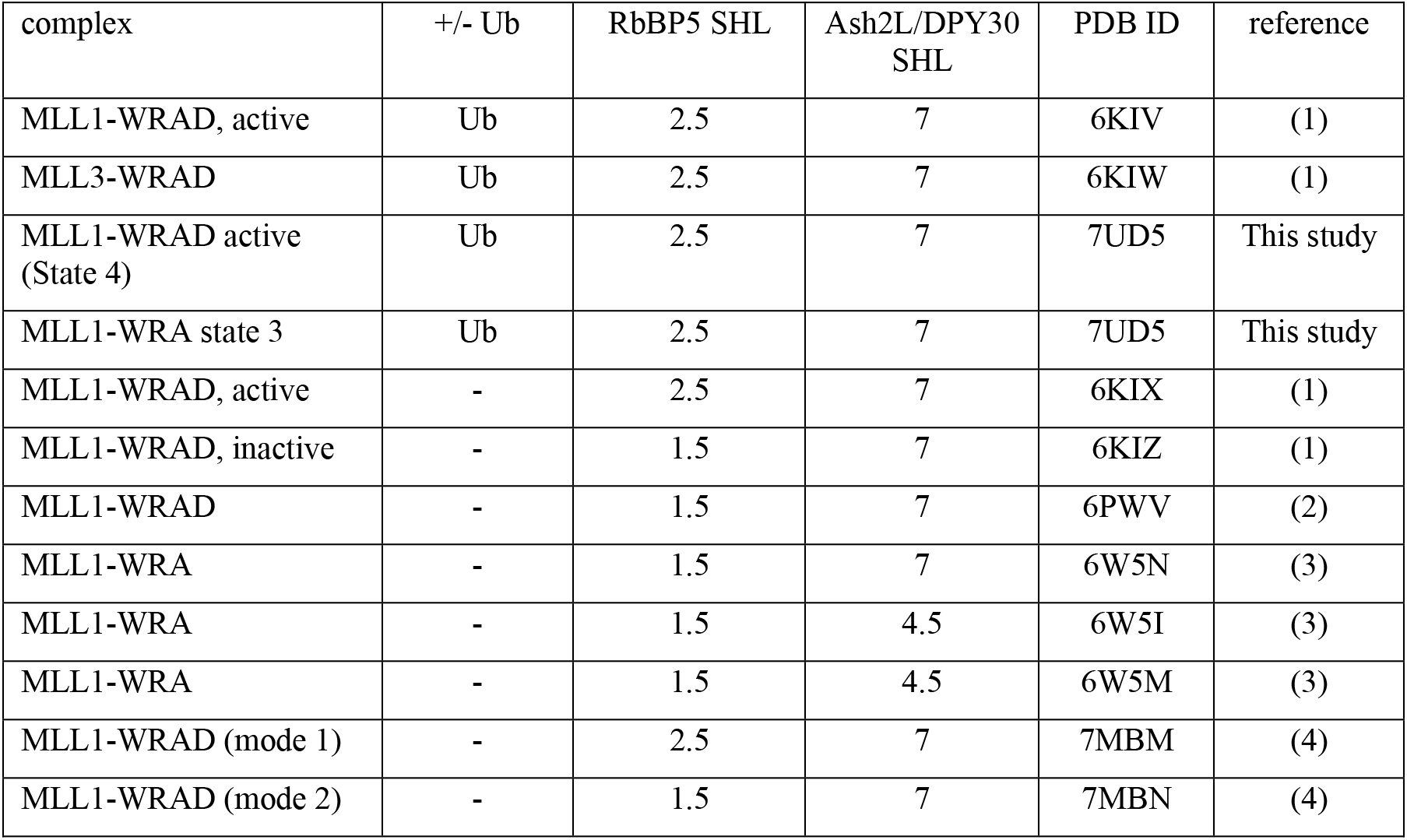
Positioning of MLL complexes on the nucleosome.

A comparison of structures determined with nucleosomes containing H2B-K120Ub suggests a role for ubiquitin in positioning the MLL1-WRAD complex on the face of the nucleosome. Monoubiquitination of H2B has been shown to increase the catalytic efficiency of MLL1-WRAD on nucleosomes by 2-fold in vitro (1). The structure reported here of the active state of the MLL1-WRAD complex (Figures 1b and 6d) is docked on the H2B-ubiquitinated nucleosome in essentially the same orientation as a previously reported structure of MLL1-WRAD bound to H2B-ubiquitinated nucleosome (6KIV, (1)), with RbBP5 at SHL 2.5 and Ash2L at SHL7. The MLL3-WRAD complex, which contains a related methyltransferase subunit and the identical four adapter proteins, is oriented in a similar manner on H2B-ubiquitinated nucleosome (6KIW, (1)). By contrast, structures of MLL1 complexes bound to unmodified nucleosomes show highly variable positioning of the MLL1 complex. Two reported structures of MLL1-WRAD bound to unmodified nucleosomes, 6PWV (2) and 6KIZ (1), also have the Ash2L subunit positioned around SHL 7, but with the complex markedly pivoted by about 35° (Table 1 and Supplementary Figure S5A and S5B). This rotation places RbBP5 in contact with nucleosomal DNA at SHL 1.5, in addition to reducing the contact area with the nucleosome. A subset of these particles adopted the active orientation (6KIW), which suggested that MLL1-WRAD could sample both positions in the absence of ubiquitin (1). A similar heterogeneity in positions resembling the active and peripheral complexes was also observed in a recent study of MLL1-WRAD bound to unmodified nucleosomes (7MBM, 7MBN, (55)). Taken together, the strong correlation of H2B-K120 ubiquitination with positioning of RbBP5 at SHL 2.5 and the absence of alternative docking modes suggests that the interaction with ubiquitin favors this docking arrangement, which we term the active complex.

The absence of DPY30 gives rise to further heterogeneity in docking of MLL1-complexes on nucleosomes that are not modified with ubiquitin. Structural studies of a four-protein complex lacking both DPY30 and the Ash2L SDI helix was found to bind to unmodified nucleosomes in multiple orientations (3) (6W5I, 6W5M, 6W5N; Table 1). While one class (6W5N) resembles that of the five-protein MLL1-WRAD complex bound to unmodified nucleosomes (6PWV), with Ash2L at SHL7, the remaining classes bind in another orientation, with Ash2L at SHL 4.5 (6W5I, 6W5M) (Supplmentary Figure S5C). This observation is consistent with a role for DPY30 in favoring binding between SHL6 and SHL7, as observed in structures of the complete MLL1-WRAD complex containing DPY30 (6KIV, 6KIX, 6KIZ, 6PWV and this work). DPY30 likely exerts its effect through its interactions with the Ash2L IDR, which undergoes conformational changes in the presence of DPY30 as reflected in changes in NMR spectra (3). The proximity of the Ash2L IDR and SPRY-insertion region to nucleosomal DNA in our model (Figure 5), as well as in a previous proposed model(2), could account for the coupling of DPY30 binding to ordering of this region of Ash2L.

## Discussion

Structural studies of the MLL1-WRAD complex have revealed surprising heterogeneity in the positioning of the complex on nucleosomes. Since the substrate lysine, K4, lies in the flexible tail of histone H3 and can therefore access the active site of MLL1 in all reported structures, there has been some question as to which one represents the active conformation (2,55). The structure of the MLL1-WRAD complex bound to an H2B-ubiquitinated nucleosome reported by Xue et al. (1) most closely resembles the docking of yeast homologue, COMPASS, on nucleosomes (52,56) and was, in part, for this reason referred to as the active complex. However, in contrast with the intrinsic mobility of MLL1-WRAD, structures of COMPASS revealed a stable nucleosome-bound complex with differences in docking between ubiquitinated and unmodified nucleosomes (52,53). Unlike COMPASS, structures of MLL1-WRAD bound to unmodified nucleosomes have revealed multiple docking arrangements (1-3,55) (Table 1). The structure of the MLL1-WRAD complex we report in this work, which contains an H2B-ubiquitinated nucleosome, also contains a docking arrangement that is very similar to that observed for the only previous MLL1 complex structure that also contains a ubiquitinated nucleosome (1). The same docking was also observed for the MLL3-WRAD complex bound to ubiquitinated nucleosome (1). Taken together, these observations strongly suggest that the presence of ubiquitinated H2B-K120 orients the MLL1-WRAD complex on the nucleosome and that the resulting docking represents the active conformation. This effect is mediated through contacts between ubiquitin and RbBP5, which positions this subunit at SHL 2.5 (Table 1). In the absence of ubiquitin, the full MLL1-WRAD complex binds in a markedly different orientation that places RbBP5 at SHL1.5 and also had reduced contacts with the face of the histone octamer (2). Since H2B-ubiquitination increases methyltransferase activity in vitro by about 2-fold (1), it is likely that the docking observed in the absence of ubiquitination simply represents a lower activity state, rather than an inactive state. H2B-ubiquitination could enhance activity by favoring docking of MLL1 in the more active conformation.

The role of H2B-K120Ub in positioning MLL1 in an active docking orientation most closely resembles the mechanism by which H2B-K120Ub activates the histone H379 methyltransferase, Dot1L. Whereas Dot1L also forms higher order complexes with partner proteins such as AF4 and AF9 (57), the catalytic domain alone is strongly stimulated by the presence of monoubiquitin conjugated to histone H2B-K120 (58). Like MLL1-WRAD, H2B-ubiquitin orients Dot1L on the nucleosome via contacts with ubiquitin (46,59,60), but in the absence of ubiquitin, Dot1L can adopt other, inactive docking arrangements (60,61). Since the H3K79 side chain is in the globular core of the nucleosome and not, like H3K4, in a flexible tail, correct positioning of Dot1L is all the more critical to its activity. In the case of Dot1L, however, ubiquitin positions the C-terminal portion of the catalytic domain but still allows Dot1L to pivot between a poised orientation, in which the active site is ∼20 Å from the substrate lysine, and an active position in which Dot1L also induces a conformational change in histone H3 (46). It is not known what role, if any, Dot1L partner proteins may play in further favoring the full active conformation on ubiquitinated nucleosomes.

Our study also points to a role for Ash2L in defining a second docking point for the MLL1-WRAD complex on the nucleosome. By combining cryo-EM data with MS-crosslinking and MDFF-refinement, we have been able to propose a new model for the Ash2L IDR (Figure 4) that differs from a previous model in which the MLL1 complex is positioned on the nucleosome in the low activity conformation (2). We also propose a model for docking of the Ash2L WH-PHD domain based on cross-linking data (Figure 5). Since our model of Ash2L predicts contacts with the DNA only, the similar relative position of Ash2L in both the active and inactive state suggests that there might be additional contacts with the histone core that further orient Ash2L.

We have identified three additional substates of the MLL1-WRAD complex that share the same overall positioning as the full active state but lack one or more subunits (Figure 6). An interesting question is whether the four states represent assembly intermediates of the active, methylating complex. Biochemical evidence suggests that H3K4 methylation requires association of the MLL1-SET domain with the nucleosome octamer (62,63), even though H3K4 lies in the flexible N-terminal tail of histone H3. The position of the MLL1-SET domain on the histone octamer surface in our active state structure agrees well with the histone residues that were identified as important to H3K4 methylation by the yeast MLL1 complex homologue, COMPASS (62). We speculate that the three less complete MLL1-WRAD states on the nucleosome, states 1, 2 and 3, may represent intermediates in assembly of MLL1-WRAD on the nucleosome. The stable RbBP5-WD40 domain present in all 4 states could in fact be an anchor for the complex that then allows further stabilization on the nucleosome. An alternative possibility is that the remaining subunits of the MLL-1 complex are flexibly tethered, and that the different states observed represent stepwise tethering of the full complex on the nucleosome surface. The significance and roles of the alternative states observed will require further study.

## Acknowledgements

We thank Duncan Sousa for assistance with data collection. We also thank Dr. Phil Gafkin at the Fred Hutchison Cancer Research Center proteomic facility, and Dr. Ebbing De Jong at the SUNY-Upstate proteomics facility for their help with mass spectrometry. Computational resources were provided by the Maryland Advanced Research Computing Center (MARCC).

## MATERIALS AND METHODS

### Expression and purification of the MLL1 complex

The MLL1-WRAD complex was reconstituted from proteins expressed in *E. coli* (21,66).

A human MLL1 construct comprising residues 3745–3969, as well as full-length human WDR5, RbBP5, and ASH2L proteins were individually expressed in *Escherichia coli* (Rosetta II, Novagen) and purified as described previously (66). DPY30 was expressed and reconstituted with the other four proteins as described in (21).

### Preparation of ubiquitinated nucleosomes containing norleucine

Nucleosome core particles containing histone H3 with norleucine in place of lysine 4 as well as H2B ubiquitinated at K120 via a non-hydrolyzable dichloroacetone (DCA) linkage were prepared as described(52). Briefly, histones H2A, H4 and the Widom 601 DNA were expressed and purified as described in (67). Histone H3 containing norleucine in place of lysine 4 was expressed and purified as described in (46). Ubiquitinated H2B was generated by expressing and purifying H2B K120C and ubiquitin K79C, which were cross-linked with DCA as described in (44). A complete protocol for generating ubiquitinated nucleosomes can be found in (68).

### Cross-linking mass-spectrometry

MLL1-WRAD complex was buffer exchanged into a crosslinking compatible buffer (50 mM HEPES pH 8.0, 200mM KCl, 5 mM MgCl2, 0.1 mM EDTA, 5% Glycerol, 1 mM TCEP). 20 ug of the buffer exchanged complex was crosslinked with 2 mM and 4 mM BS3 for 4 hours at 4°C and frozen at −20°C until further processing for LC-MS/MS as described in (69) and (70) with minor modifications. Briefly, crosslinked complexes were thawed at room temperature and an equal volume of trifluoroethanol was added and incubated at 60°C for 30 min to denature. The proteins were then reduced by addition of 5 mM TCEP for 30 min at 37°C followed by alkylation with iodoacetamide at a 10-mM final concentration for 30 min in the dark at room temperature. The sample was diluted 10-fold with 20 mM triethanolamine, and then digested with 2 μg of trypsin (Promega) overnight at 37°C. The peptides were further purified on Sep-Pak C18 cartridge (Waters), dried and resuspended in 5% acetonitrile/0.1% TFA solution and then analyzed by LC-MS/MS. BS3–cross-linked peptides were analyzed on a Thermo Scientific Orbitrap Elite and Lumos with HCD fragmentation and serial MS events that included one FTMS1 event at 30,000 resolution followed by ten FTMS2 events at 15,000 resolution. Other instrument settings included: MS mass range greater than 1,800; m/z value as masses enabled; charge-state rejection: +1, +2, and unassigned charges; monoisotopic precursor selection enabled; dynamic exclusion enabled: repeat count 1, exclusion list size 500, exclusion duration 30 s; HCD normalized collision energy 35%, isolation width 3 Da, minimum signal count 5,000; and FTMS MSn AGC target 50,000. The RAW files were converted to mgf files and analyzed by the cross-link database–searching algorithm pLink2 under default settings: (i) up to three missed cleavages; (ii) differential oxidation modification on methionine (+15.9949 Da); (iii) differential modification on the peptide N-terminal glutamate residues (−18.0106 Da) or N-terminal glutamine residues (−17.0265 Da); (iv) static modification on cysteines (+57.0215 Da); (v) 2% FDR. All possible tryptic peptide pairs within 20 p.p.m. of the precursor mass are used for cross-linked peptide searches. The cross-linked peptides were considered confidently identified if at least four consecutive b or y ions for each peptide were observed. Cross-links used for this study are in Supplementary Table S2.

### Sample preparation for cryo EM

To form stable MLL1-WRAD-nucleosome complexes, 100 nM of modified nucleosome was mixed with 5x WRAD in EM-buffer (20 mM HEPES, 300 mM NaCl, 2 mM TCEP, 0.1µM Zinc chloride) and incubated for 1 h on ice. The sample mix was then crosslinked using fresh 0.05% (w/v) glutaraldehyde for 1 h on ice and quenched with 100 mM Tris for 2 hours. The sample was diluted to 0.5 mg/ml and directly applied to freshly glow-discharged 2/2 Cu Quantifoil grids in a Vitrobot Mark IV (Thermo Fisher Scientific) at 4° C and 100% humidity. The sample was immediately blotted (3 s blot time) and flash-frozen in liquid ethane.

### Cryo EM data acquisition and image processing

All data were acquired at the Beckman Cryo EM Center (Johns Hopkins) on a Thermo Fisher Titan Krios G3 equipped with a Gatan K3 direct electron detector at a magnification of 21,500 in super-resolution counting mode, corresponding to a pixel-size of 0.529 Å/pix. A total of 5091 movies were recorded using Serial-EM (71) using a varying negative defocus of 1.0 – 2.8 µm and recording 40 frames at 1.5 e/Å^2^/frame at 60 e/Å^2^ total dose. Data acquisition was monitored and frequently evaluated using cisTEM (72). Movie stacks were aligned and down-scaled to a pixel size of 1.058 Å/pix (bin 1) using MotionCor2 (73) and CTF correction was performed using Ctffind4 (74). The full dataset was then manually inspected and 4804 movie stacks were selected for further processing in Relion 3.0 (75). Initial particle picking yielded 2,511,854 particles that were subjected to 2D and 3D classification using 3-fold downscaled particle images (bin3) for faster processing speed and stronger signal-to-noise ratio, which removed junk particles and yielded 608,972 ‘good’ particles that displayed MLL1-complex bound to the nucleosome. All ‘good’ particles were re-extracted at 1.058 Å/pix (bin1), consensus-refined, and subjected to masked 3D-classification using a Ash2L-DNA interface mask, which yielded six unique classes displaying differences in Ash2L density and DNA topology (see Extended Data Figure S1). The 103,849 particles were subjected to one round of CTF refinement followed by Bayesian polishing as implemented in Relion-3.0. This procedure yielded a reconstruction after refinement of 3.33 Å resolution based on the FSC 0.143 (76) criterion termed State 1 (see Extended Data Figure S1, sS2), which was sharpened using an automatically calculated B-factor of −70 Å^2^. Particles grouped in classes 2 and 5 from the initial masked classification were combined after confirming similar classes by visual inspection and reclassified into four classes (see Extended Data Figure S1). The resulting best class containing 134,528 particles showed flexible Ash2L and MLL1 density and density stretches associated to the DNA between SHL −6 to −7 and was termed State 2. Particles were subjected to one round of CTF-refinement followed by Bayesian polishing, yielding a 3.55 Å resolution reconstruction (according to the FSC 0.143 criterion) that was sharpened using an automatically calculated B-factor of −86Å^2^. The 56,526 particles grouped in class 3 from the initial masked classification were CTF-corrected, particle polished, and subsequently refined to a final resolution of 4.65 Å (0.143 criterion). This reconstruction and sharpened using an automatically calculated B-factor of −117 Å^2^ (see Extended Data Figure S2) and termed State 3. The 66,449 particles grouped in class 4 from the initial masked classification were CTF-refined, particle polished and further refined to a final resolution of 3.95 Å based on the 0.143 criterion). This class contained highly resolved nucleosome density but more poorly resolved MLL1 complex and Ash2L density. In order to further deblur the Ash2L components in this state, an additional 3D-classification using only local searches was performed (see Extended Data Figure S1). This process yielded an overall better-resolved class (25,525 particles) that refined to 4.25 Å after postprocessing and was sharpened using a B-factor of −75Å^2^. This map (State 4) was used to guide Ash2L modelling as described below.

### Model building and refinement

Crystal structures of human ubiquitin (1UBQ), human WDR5^31-334^ (3EG6), human MLL1^3814-3969^-Ash2L^285-504^-RbBP5^330-375^ (5F6L), DPY30^50-96^ (6E2H), and *X. laevis* nucleosome (6KIU) were subjected to rigid body fitting in the State 4 map using Chimera (50). These combined fitted structures allowed to allocate most resolved density of the State 4 map except a stretch of continuous density (see Figure 1c) presumably corresponding to Ash2L based on homology of the complex to solved structures from other organisms (see Figure 3b and c). The Ash2L model was computed through iterative rounds of crosslinking-based homology modelling utilizing available structures and manual model-building, followed by Molecular Dynamics Flexible Fitting (MDFF) using our State 4 map. Specifically, we use the previously resolved SPRY domain (287-504) from the MLL1^SET^-ASH2L^SPRY^-RbBP5^330-375^ complex (PDB ID: 5F6L) (40) and added residues 398-440 from a recently reported Ash2L homology model (2). All MDFF simulations were done in NAMD2.13 (77). The CHARMM36 force field (78,79) was used for both protein and DNA. All simulations were carried out at 300 K in vacuum with a scaling factor of 1.0. To avoid potential structural artifacts that can arise from MDFF, secondary structure, chirality, and cis-peptide restraints were applied. For targeted refinement of Ash2L and DPY30, a 500 kcal/mol/Å harmonic restraint was applied on all backbone atoms except for Ash2L and DPY30. The cross-correlation coefficient was computed using the MDFF package implemented in VMD 1.9.3 (80).

Next, we manually optimized the models fit to our experimental density, which generated our initial Ash2L-model. We assigned adjacent helical tube density to DPY30 and performed automated fitting of the Ash2L-SDI / DPY30 dimer crystal structure (PDB: 6E2H). In addition, our density allowed us to fully build the C-terminal helix of Ash2L (487-528). Ash2L residues 199-286 were built using MODELLER (81) using cross-link restraints and underwent iterative rounds of manual model-building guided by the EM density. The model was further refined through a 3.0 ns MDFF simulation (43). To improve the overall fit of Ash2L and DPY30, the 6.0 Å active state map (Ash2L-focused) was used as the template map. After refinement with MDFF, the cross-correlation coefficient between the overall structure and template map improved from 0.79 to 0.83. To correct any rotamer outliers that may occur from MDFF refinement, the model of the full complex underwent 1000 iterations of minimization with secondary structure restraints using the Phenix geometry minimization module. Following, the model was iteratively refined with Phenix real-space refinement. Final statistics for the State 4 model are shown in Supplementary Table S1.

### Ash2L N-terminal Docking

Cross-link guided docking of the human Ash2L PHD-WH domain (PDB: 3RSN) was carried out with HADDOCK 2.4 (82). Crosslinks were used as unambiguous restraints between the Ash2L N-terminal residues and our reported MLL1-WRAD complex. In HADDOCK, 5000 rigid-body calculations were first carried out and the top 1000 structures underwent semiflexible refinement. Finally, HADDOCK produced 200 water-refined structures. Clusters were defined using Fraction of Common Contacts (FCC) with a cutoff of 0.60. All 200 structures were identified to a single cluster with a final HADDOCK score of −213.7±1.4.

## Data availability

Coordinates of the full active complex (state 1) have been deposited in the Protein Data Bank, PDB ID 7UD5. Maps have been deposited in the Electron Microscopy Data Bank, EMD-26454.

## Supplementary figures

**Figure S1.**
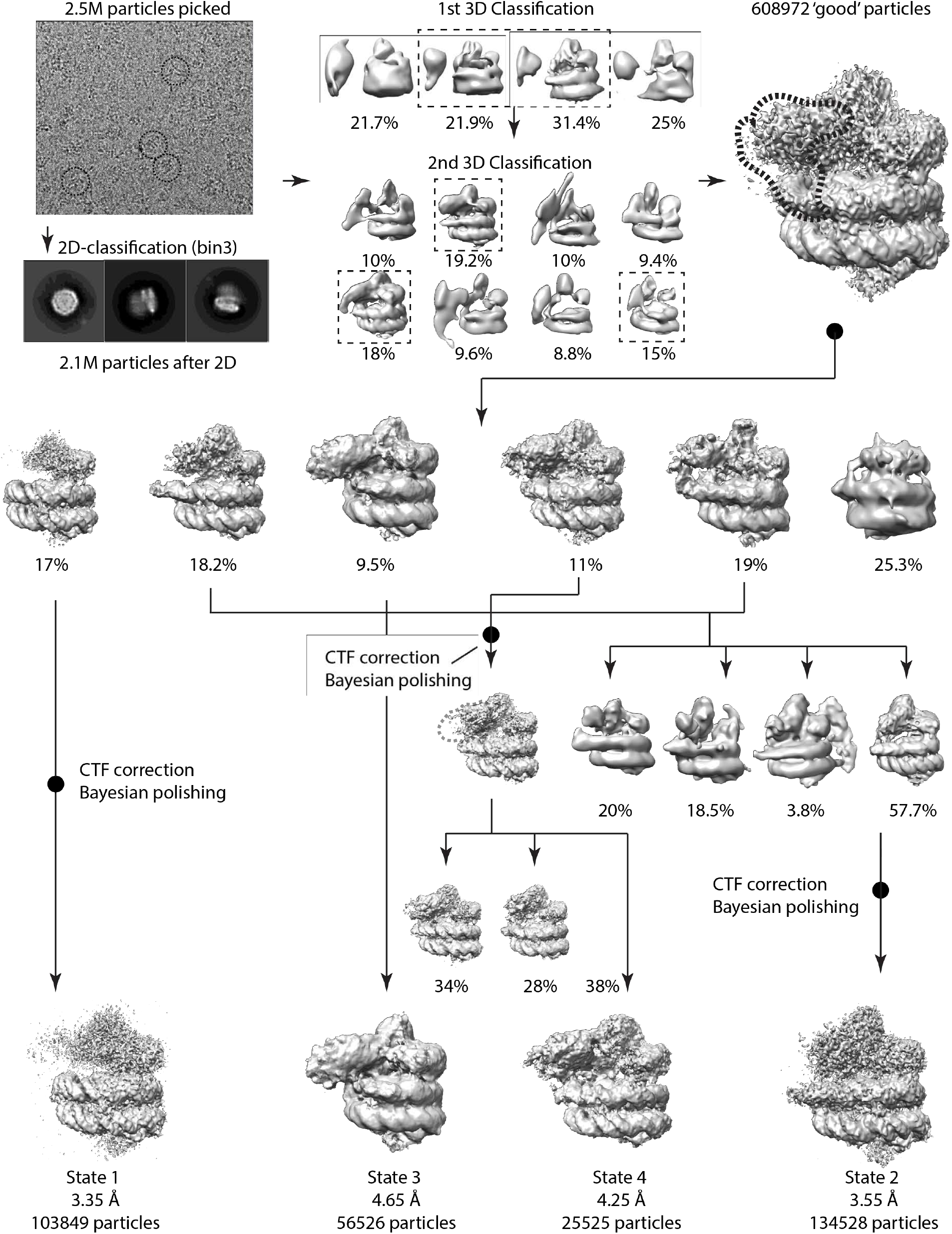
Processing pipeline depicting steps and intermediate states. All maps are displayed after refinement.

**Figure S2.**
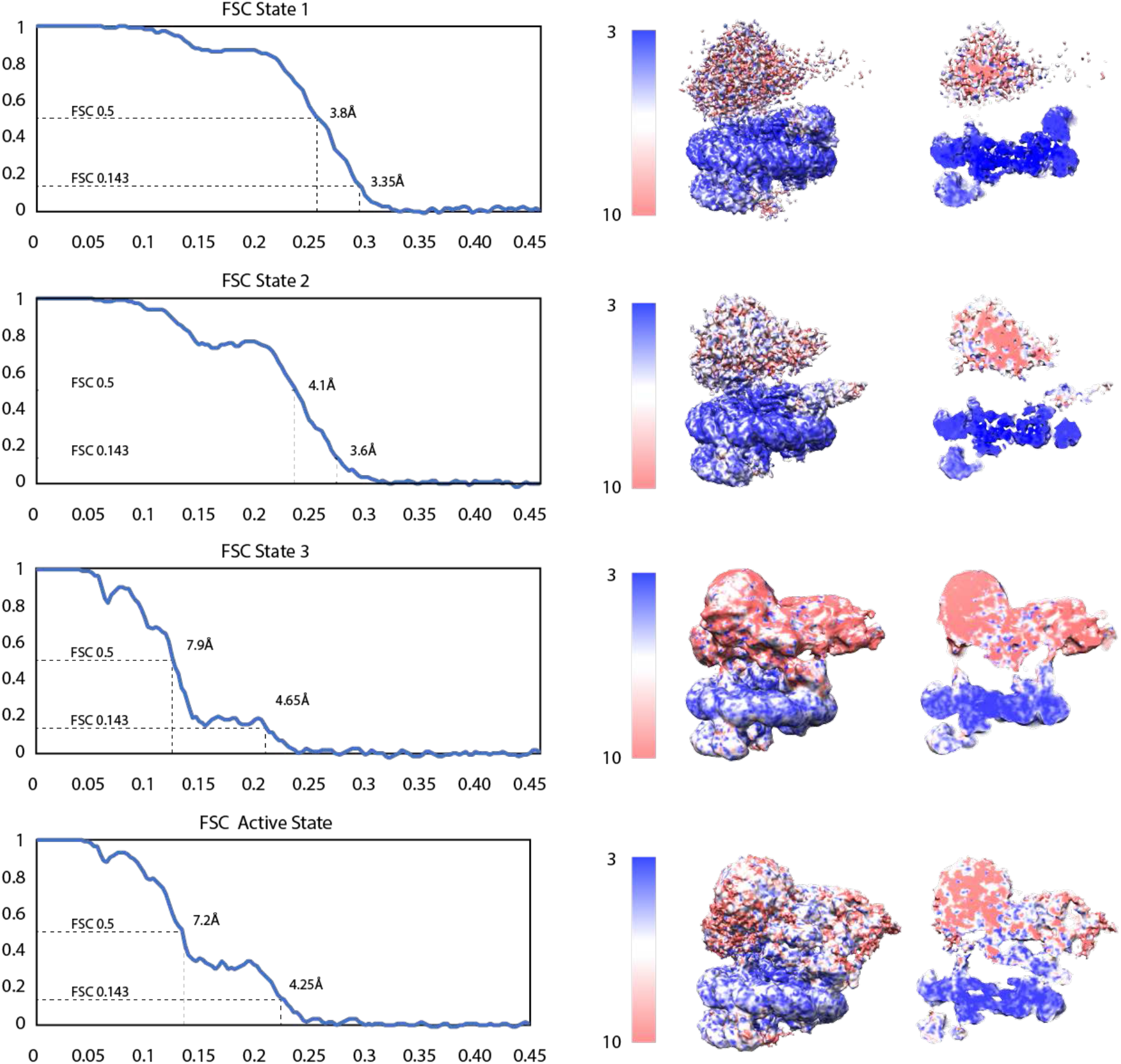
Fourier Shell Correlation and Local Resolution for the four MLL1-WRAD states. For each state, Fourier shell correlation (FSC) curves (left panel) and associated local resolution(right panel) are displayed.

**Figure S3.**
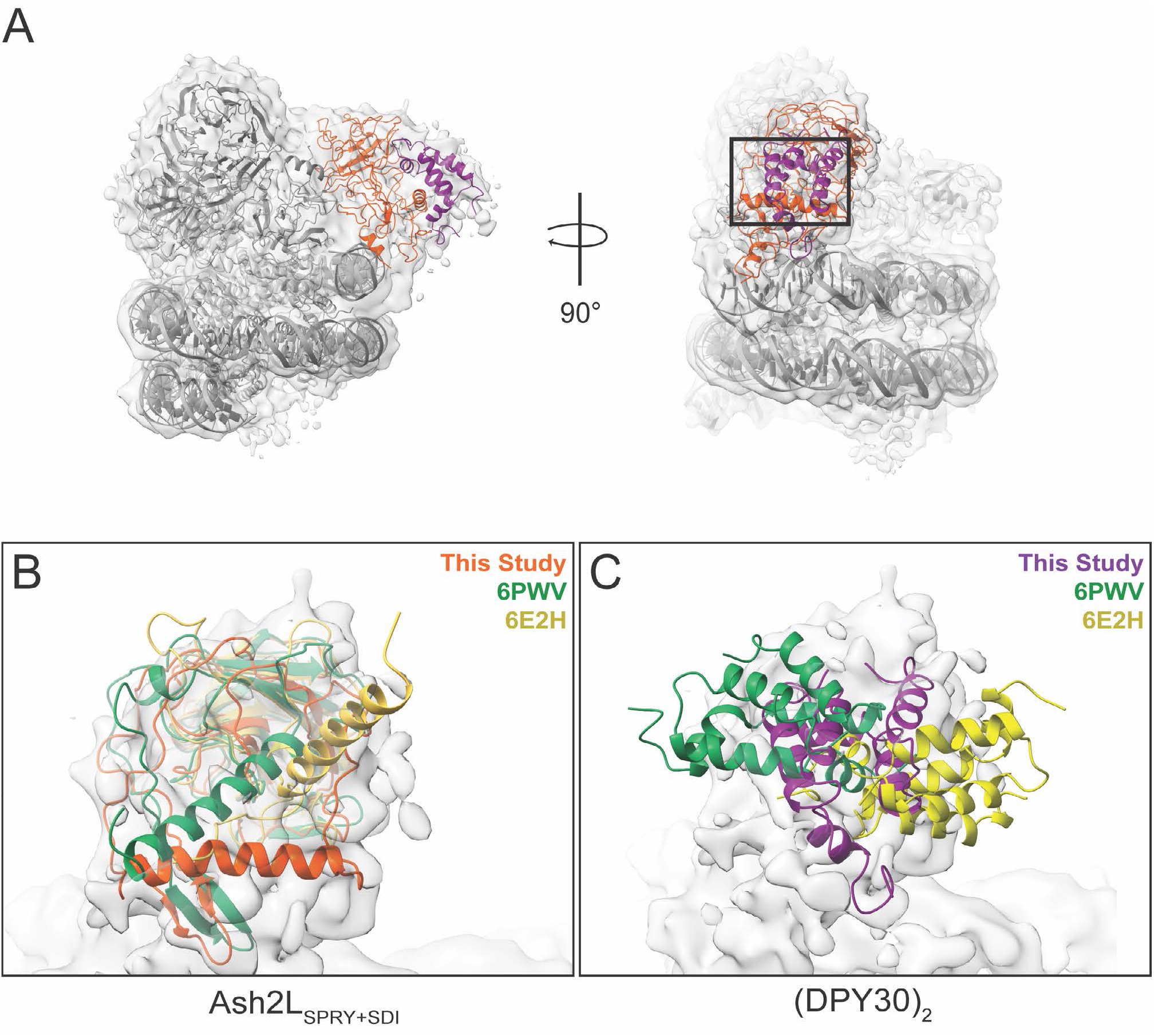
Comparison of reported structures of the Ash2L-SDI motif and DPY30 dimers against active state map. A) Our model of Ash2L and DPY30 fit into the active state map. B) Ash2L-SDI motif of the model presented in this study (orange), Park et al. (green; 6PWV), and Haddad et al. (yellow; 6E2H). Across the three models, the SDI helix vary in orientation. However, only the SDI-motif of our model and Park et al. fits well in our active state map. C) DPY30 dimer position reported in this study (purple), Park et al. (green; 6PWV), and Haddad et al. (yellow; 6E2H). The orientation of DPY30 reported by Park et al. and Haddad et al. fit poorly into our active state map. Differences in orientation of DPY30 is attributed to the differences in the SDI motif orientation across the three models.

**Figure S4.**
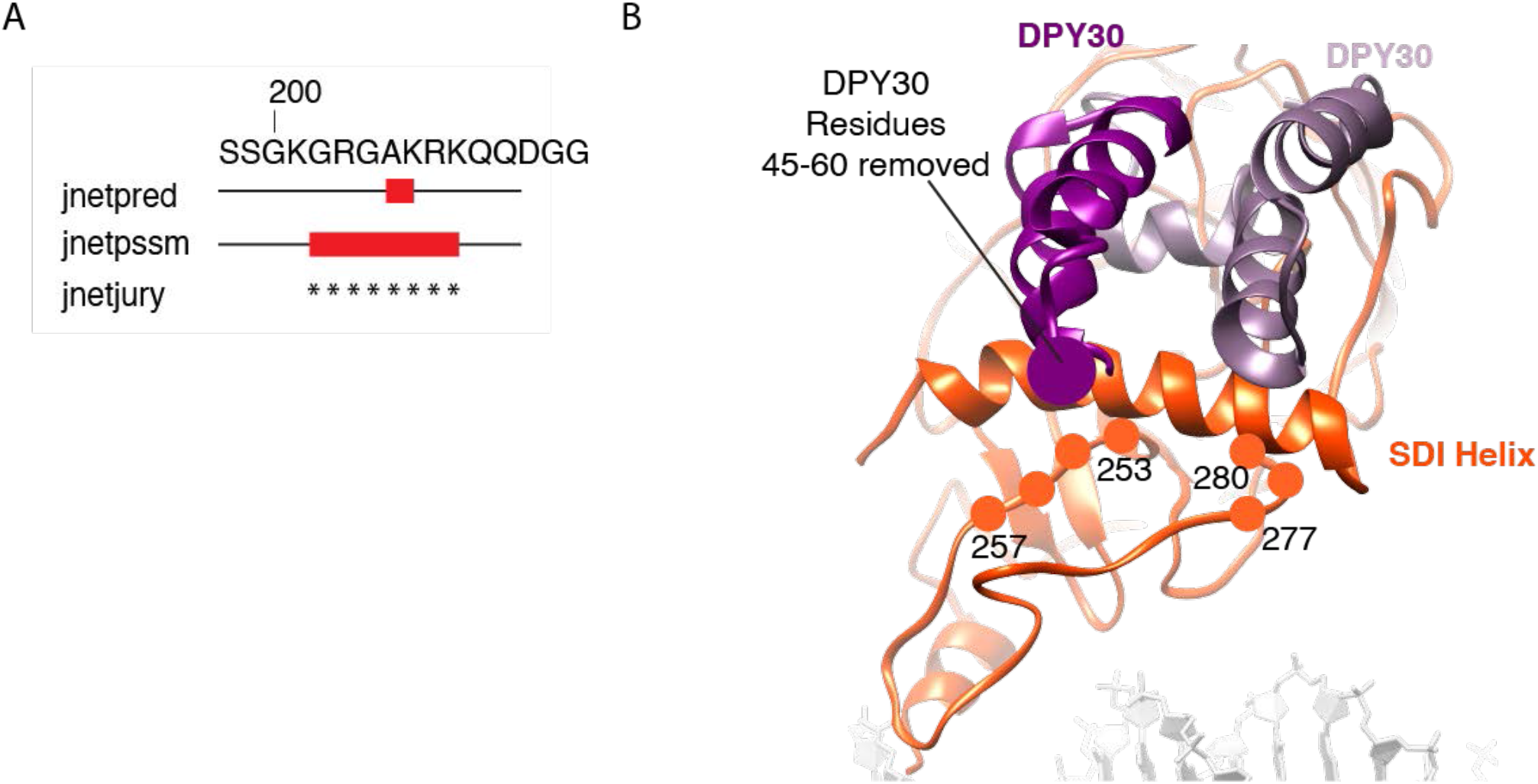
Modeling of the Ash2L intrinsically disordered region (IDR) A) Secondary structure prediction of alpha-helix spanning Ash2L residues 202-209. B) Ash2L-SDI helix and associated IDR-stretch. IDR residues 253-257 and 277-280 are depicted as balls and have been shown to impede H3K4 methylation when mutated to alanines.

**Figure S5.**
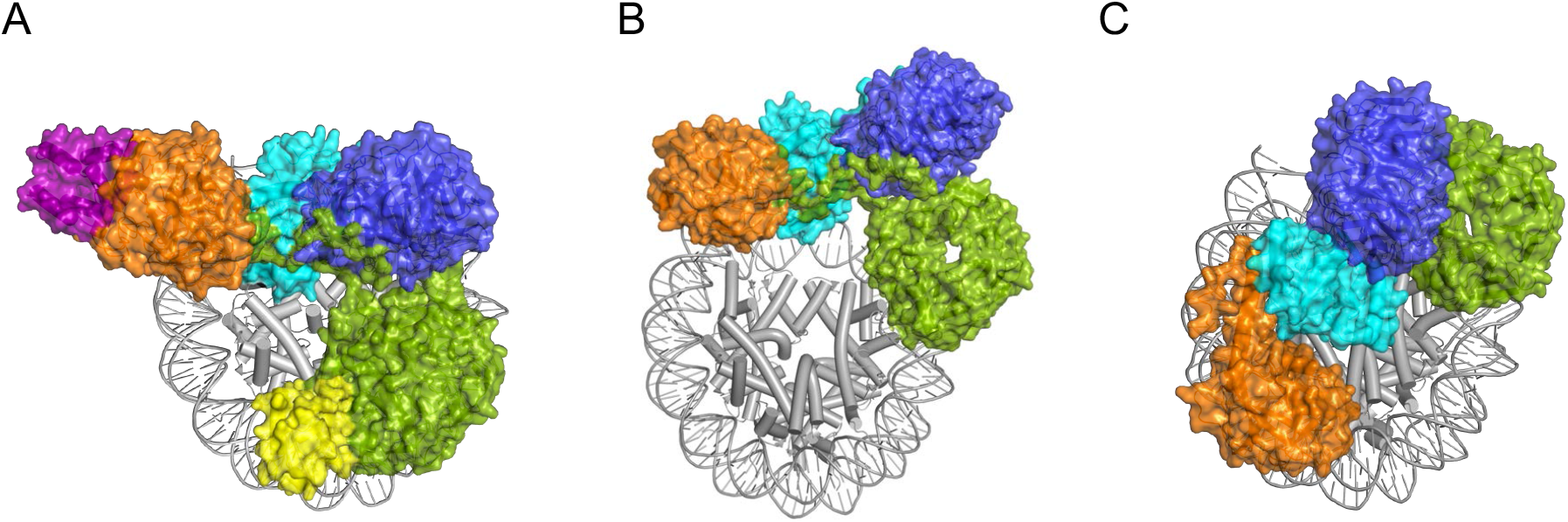
Position of MLL1 complexes on the nucleosomes. Color scheme for all panels; ubiquitin-yellow, RbBP5-green, WDR5-blue, MLL1-cyan, Ash2L – orange, DPY30 – purple. A) MLL1-WRAD complex bound to H2B-ubiquitinated nucleosome, present study (7UD5). B) MLL1-WRAD bound to unmodified nucleosomes (6KIZ). C) MLL-WRA bound to unmodified nucleosome (6W5M).

**Table S1:**
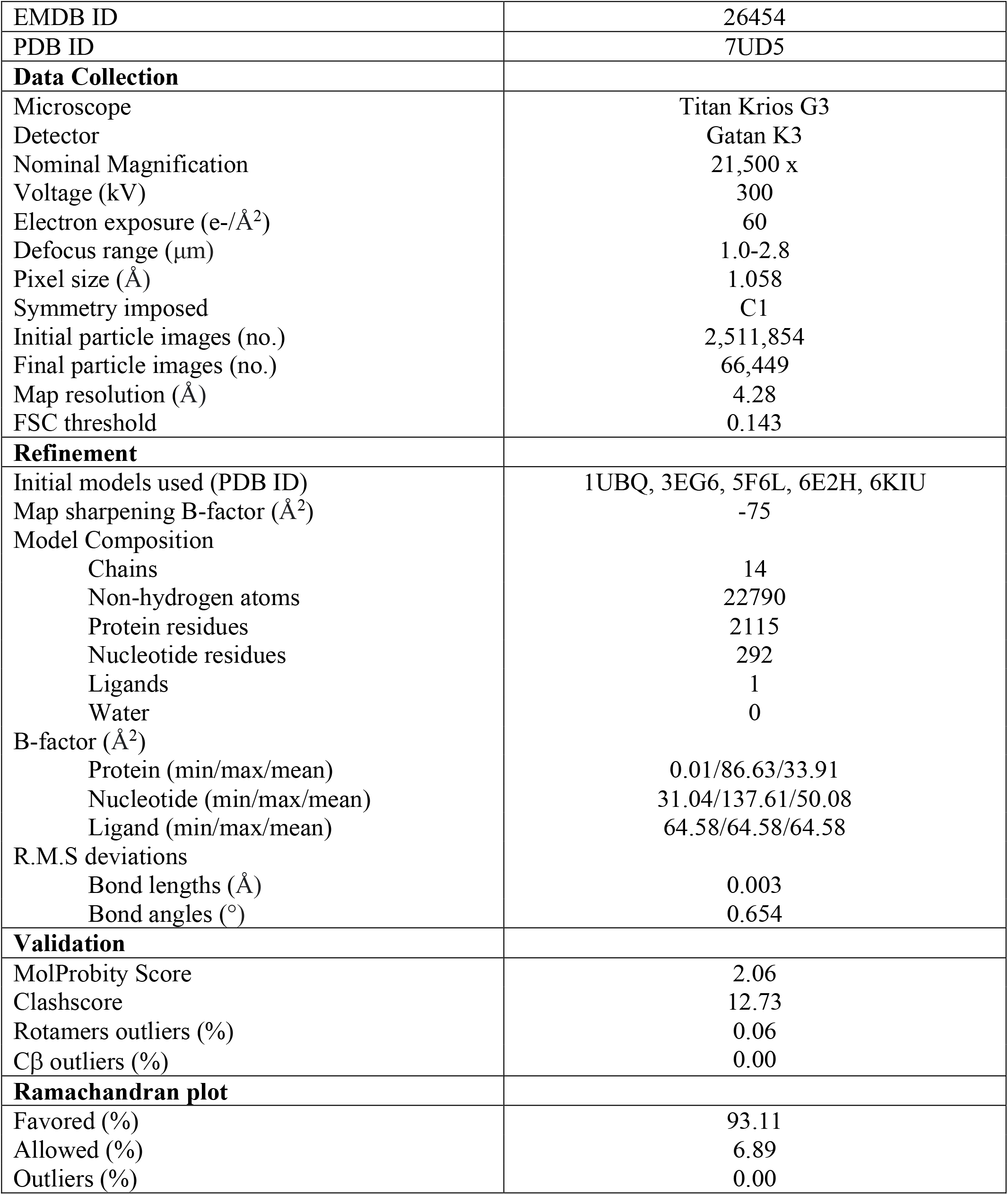
Cryo-EM data collection, refinement, and validation statistics.

**Table S2:**
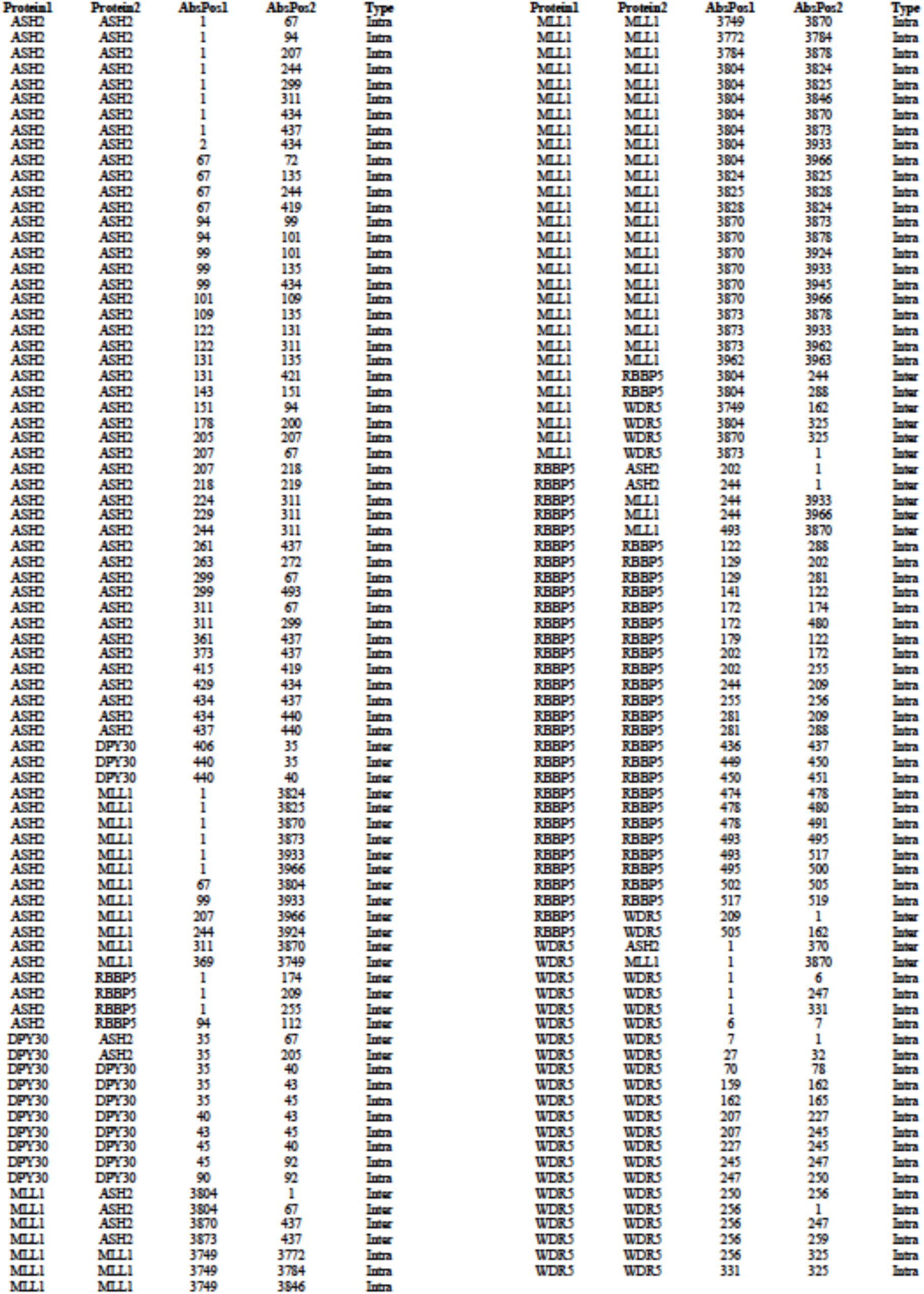
Recorded Crosslinks (2% false discovery rate)

